# Early excitatory and inhibitory modifications in the motor cortex following skill learning support motor memory consolidation and cortical plasticity overnight

**DOI:** 10.1101/2022.12.27.521981

**Authors:** Tamir Eisenstein, Edna Furman-Haran, Assaf Tal

## Abstract

The learning of new motor skills constitutes an inseparable part of our lives. Motor consolidation refers to the offline processing of motor memories following the acquisition of new motor skills. The animal literature suggests that the primary motor cortex (M1) plays a key role in motor memory consolidation, and structural and functional plasticity in M1 following motor consolidation have been demonstrated. However, the mechanisms supporting motor memory consolidation and plasticity in the human M1 are not well understood. Initial human neuroimaging studies show that the initial stages of motor learning in humans are accompanied by short-term temporal dynamics of the brain’s main excitatory and inhibitory neurotransmitters – Glutamate (Glu) and GABA – in M1, but it remains unclear how these relate to the question of motor memory consolidation.

Here, we show that early Glu and GABA modifications in M1 following motor skill learning may play vital roles in supporting motor memory consolidation and neural plasticity that take place over longer time scales. Using a multimodal magnetic resonance approach implemented on ultra-high field 7T scanner in healthy young adults (n=36), we found increased Glu and decreased GABA in M1 during the initial offline period following learning to support consolidation-related local and inter-regional functions of M1, such as motor memory reactivation and increased functional connectivity with the striatum. These neurochemical changes also correlated with overnight structural and functional plasticity expressed as increased M1 grey matter volume and functional connectivity, while Glu modifications also correlated with adaptive behavior, as reflected by improvements in skill performance.

Our results provide intriguing microscale mechanistic evidence to the potential distinctive roles of Glu and GABA in promoting motor memory consolidation and plasticity in the human M1. They also highlight a role for early neurochemical modifications to memory consolidation and plasticity in the human brain and may hold important clinical implications in rehabilitative settings such as in stroke and brain injury.

## 1. Introduction

The learning of new motor skills constitutes an inseparable part of our lives. Acquiring new motor skills is manifested from new-borns learning to grasp a spoon, through learning a new piece on the piano, and to the re-learning of basic motor skills following brain injury. Motor skill learning is a gradual continuous process that is characterized by an initial stage in which performance increases relatively fast (i.e., early phase), followed by a later slower stage in which further gains in performance develop gradually over additional practice (i.e., late phase)^1^. The duration of each stage is highly dependent on the complexity of the acquired motor skill, and both stages may be evident on the first practice session when learning simple skills such as short key-press sequences^2,3^.

Following their initial encoding/acquisition, motor memories are continued to be processed in the brain, in an offline process termed consolidation. During consolidation, the motor memory is thought to be stabilized, strengthened, and re-organized in the brain^4^. Previous studies have highlighted the importance of the first hours following learning to the stabilization of the motor memory, while the offline enhancement of skill performance usually develops across sleep ^5,6^. However, despite the obvious importance of consolidation to motor skill learning, the neural mechanisms underlying this process are not entirely clear.

The primary motor cortex (M1) plays a crucial role in both the acquisition and consolidation of new motor skills, which in turn result in the formation of a stable neural representation of the acquired skill in the brain, or a motor memory “engram”^7^. Furthermore, motor memory consolidation has been shown to be associated with structural and functional plasticity in M1 on multiple levels^1^. Microstructural changes, such as dendritic spines remodelling, have been demonstrated in animals following motor learning^7^, while macroscopic changes in grey matter (GM) volume were observed in humans^8^. In addition, changes in the functional communication between M1 and related brain regions have been shown following the acquisition of new skills^9–12^, highlighting the importance of early post-learning functional modifications to the learning outcomes ^9^. Moreover, offline motor memory reactivation, a putative mechanism of memory consolidation^13,14^, has been demonstrated in M1 in animals^15^, and evidence for motor memory reactivation has also been demonstrated in humans^3^, by demonstrating a resemblance of neural activity patterns during rest following learning to those observed during the learning experience itself^16^. However, while significant progress has been made regarding the offline macroscopic and systems-level modifications, the microscale mechanisms which may underlie motor consolidation and skill learning-induced plasticity in the human brain remain unclear.

Neural excitation and inhibition (E-I) have been proposed to play an important role in the physiological regulation of cognition and behaviour, at both the single neuron level and large-scale circuits^17^. E-I are mediated by the main excitatory and inhibitory neurotransmitters glutamate (Glu) and γ-aminobutyric acid (GABA), respectively. While the physiological balance between excitation and inhibition in the brain is homeostatically regulated, dynamic responses of Glu and GABA induced by external inputs and internal processes have been suggested to play a vital role in plasticity processes and adaptive behaviour following learning^18,19^. For example, previous works in animals have shown that transient E-I alternations such as GABA-mediated cortical disinhibition support M1 plasticity ^20^. Furthermore, modulation of glutamatergic and GABAergic processing, both in vitro and following learning, has been implicated in the induction of long-term potentiation (LTP) and the promotion of synaptic strengthening^20,21^.

Magnetic resonance spectroscopy (MRS) is currently the only method capable of non-invasive examination of the molecular neurochemical environment in the human brain, including the quantification of Glu and GABA^22^. Using MRS, evidence for a role of E-I in the learning of new memories in the human brain have been proposed^11,19,23–25^. Interestingly, while motor learning was previously linked with the dynamics of *GABA* in M1^19,24^, declarative learning was related to *glutamatergic* responses in the hippocampus^25^. Yet, at present, most of the studies aiming to understand the neurochemical underpinnings of learning and memory have focused on the online phase of learning, when the experience is being encoded. As a result, we lack significant knowledge of the neurochemical responses taking place following learning^26^ and how they relate to learning-induced neuro-behavioral changes, especially during the initial and important period of offline consolidation^5,6^.

Here, we aimed to take advantage of the increased spatial, temporal, and spectral resolution of ultra-high field 7T MRS to investigate the hypothesis that early modifications in GABA and/or Glu following skill learning play important role in the consolidation of the new motor memory as reflected by overnight improvement in skill performance^5,27^. Furthermore, by also acquiring multimodal structural and BOLD MRI data, we examined how changes in Glu and GABA are associated with local and remote functional processing in M1 immediately following learning. Specifically, we used a multivoxel local correlation pattern (MVLC) analysis, similar to a previously used approach for evaluating memory reactivation in the human brain^16,28^, to determine whether reactivation may depend on transient changes in Glu and GABA following learning. Cortico-striatal interactions are critically implicated in the consolidation of motor memories^7,12,29^; We therefore also examined the relationship between the dynamics of Glu and GABA in M1 shortly after learning, and the inter-regional communication of M1 and the putamen as quantified using resting state BOLD data. Lastly, since motor skill learning has been shown to induce plastic changes in M1, we investigated whether changes in Glu and GABA following learning may be related to structural and functional plasticity in M1, as expressed by overnight changes in the GM volume and functional connectivity of this region.

## 2. Materials and methods

### 2.1 Participants

36 healthy right-handed young adults (age 27.2±3.8 years, 15 females) participated in the current study. All participants provided written informed consent, approved by the Wolfson Medical Center Helsinki Committee (Holon, Israel), and the Institutional Review Board (IRB) of the Weizmann Institute of Science, Israel. Exclusions criteria included age below 18 years or above 40 years, musicians or video gamers (past or present), any neuro-psychiatric history (including medications), and participants who did not meet the safety guidelines of the 7T scanning policy.

### 2.2 Experimental Protocol

We conducted a within-subject repeated measures experiment using a multimodal MR approach implemented on an ultra-high field 7T MRI scanner (**Figure 1A**). Participants arrived at the lab at two consecutive days. During the first day, participants underwent anatomical, single-voxel MRS and resting-state fMRI scans prior to and following a motor sequence learning task. Task-induced BOLD data was recorded during the task itself. On the second day, participants underwent anatomical, resting state fMRI and MRS only once at baseline, prior to a motor learning evaluation fMRI paradigm (i.e., a testing paradigm).

**Figure 1.**
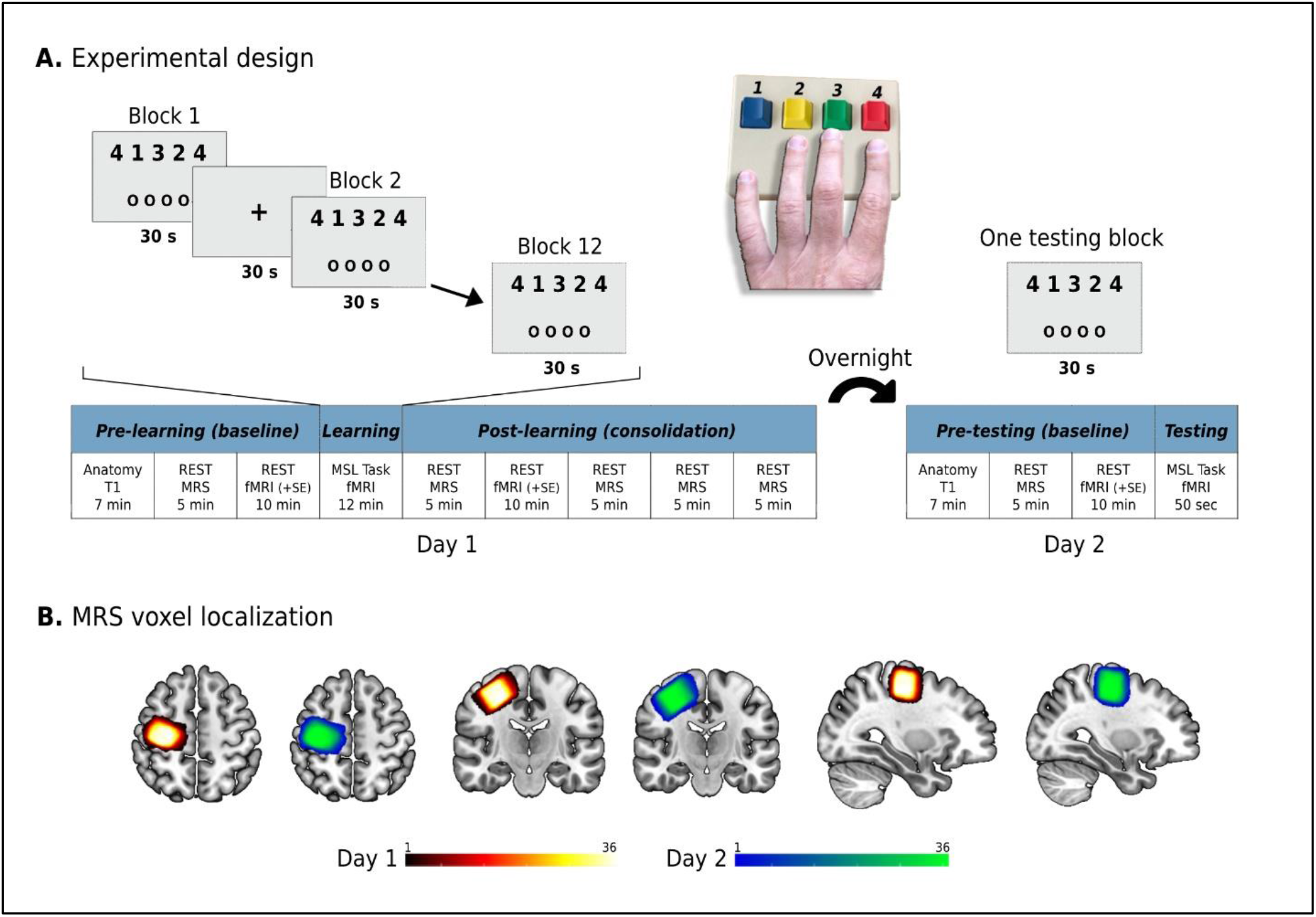
Schematic illustration of the experimental procedure of the study (**A**); MRS voxel localization during the two scanning sessions (**B**). Note the consistent voxel placement across the two experimental sessions.

### 2.3 Motor Learning Task

All participants performed an explicit motor sequence learning task in which they were asked to repetitively tap a five-digit sequence (4-1-3-2-4) with their non-dominant left hand, as fast and accurately as possible^2^. Keypresses were performed on an MR-compatible response box with four computer-like pressing keys (Cedrus Corporation, Lumina LS-LINE model). The response box was placed near the left thigh, was adjusted for each participant’s arm length, and fixed to this position to prevent its movement during the scan. Keypress 1 was performed with the little finger, keypress 2 with the ring finger, keypress 3 with the middle finger and keypress 4 with the index finger. The task consisted of 12 trials, lasting 30 seconds each during the first day (of initial learning). Each two consecutive task blocks were separated by a 30 second fixation block in which participants were asked to fixate on a black cross presented at the middle of a bright screen. The second day (of learning evaluation) consisted of a single block of 30 seconds. The five-digit sequence (4-1-3-2-4) was projected on the screen during each task block. Each task block also included four white circles presented on the screen, and each finger press was followed by a corresponding circle being filled for the time duration of that specific press. This procedure was implemented to provide the participant with online visual feedback (whether pressing on the desired key) and the experimenter with online information regarding the execution of the correct finger sequence. The visual circles did not provide error feedback, only information on the pressing finger at a given time. The first day also included a short pre-scan familiarity session with the task on a lab computer, in which only the experimenter performed a short version of the task with a control sequence for demonstration, to prevent any learning effect in the participant prior to the actual task in the scanner. Therefore, the “real” sequence was only revealed to the participants when starting the learning paradigm. Importantly, all participants demonstrated at least one correct sequence pressing during the first learning block. Behavioral performance in the task was evaluated by the number of correct sequences performed within each block, a common behavioral measure that combines both the speed and accuracy of the performed skill ^2^. We then defined the early and late phases of learning during the first day for further analyses which are described below. The late phase was defined as the practice block from which no block-to-block significant changes in performance were observed at the group-level (α<.05). This evaluation was based on non-corrected pairwise comparisons between consecutive blocks following a random-intercepts and random-slopes linear mixed model analysis with participants’ ID as the random effect. According to this, the first three blocks comprised the early phase, and blocks 4 to 12 comprised the late phase. This division is similar to what was reported in a recent study^3^ implementing the same learning sequence (when this transition occurred after ∼30% of the blocks). Visual stimuli presentation and data collection were conducted using the Psychophysics Toolbox Version 3 (http://psychtoolbox.org/) implemented in MATLAB.

### 2.4 MRI scanning procedure

#### 2.4.1 MR data acquisition

The scanning sessions were performed on a 7T Terra scanner (Siemens-Healthineers, Erlangen, Germany) using a commercial single-channel transmit/32-channel receive head coil (NOVA Medical Inc., Wilmington, MA, USA), capable of maximum B_1+_ amplitude of 25 *μ*T. Soft pads were used to hold each participant’s head in place to minimize head movement during the scanning sessions. An initial localizer and gradient echo field map were acquired for automated B_0_ shimming (Scan parameters for B0 mapping: TR/TE1/TE2=406/3.06/4.08 ms, *α*=25°, 1.9x1.9×2.0 mm resolution, TA=1:04 min). A high-resolution structural T1-weighted MP2RAGE (Magnetization Prepared 2 Rapid Acquisition Gradient Echoes) image was acquired for voxel placement and subsequent tissue segmentation (TR/TE/TI1/TI2=4460/2.19/1000/3200 ms, *α*1 = 4°/*α*2 = 4°, 1 mm^3^ isotropic voxels, TA=6:56 min). For the MRS acquisitions, a 2x2x2 cm^3^ spectroscopic voxel was placed over the hand knob region of the right primary motor cortex, based on neuroanatomical guidelines^30,31^ (**Figure 1B**). The voxel was shimmed using the automated B0 shimming capabilities of the in-house Visual Display Interface (VDI) libraries (The Weizmann Institute of Science, Israel, www.vdisoftware.net) in MATLAB 2020b (The Mathworks, Natick MA). The MRS acquisition was performed with a SemiLASER (sLASER) sequence (TR/TE=7000/80 ms, NEX=36, TA=4:58 min) previously optimized, and also validated for other brain regions as well ^32^. Functional MRI data were acquired using a multiband gradient-echo echo-planar imaging sequence. Scanning parameters were implemented according to the Human Connectome Project (HCP) 7T protocol^33^ (TR/TE=1000/22.2 ms, field of view=208×208 mm^2^, matrix size=130×130, voxel-size=1.6 mm^3^, 85 slices, multi-band/GRAPPA acceleration factor=5/2, bandwidth=1924 Hz/Px, flip angle=45°). The resting state scans included the acquisition of 420 volumes per scan, while the task paradigms included the acquisition of 710 and 50 volumes on the first and second day, respectively. In addition, spin echo field maps with opposite phase encoding directions were acquired immediately prior or following each functional acquisition for EPI distortion correction (TR/TE=3000/60 ms, field of view=208×208 mm^2^, matrix size=130×130, voxel-size=1.6 mm^3^, 85 slices, multi-band/GRAPPA acceleration factor=5/2, bandwidth=1924 Hz/Px, flip angle=180°).

#### 2.4.2 MRS analysis

MRS pre-processing was carried out using the VDI libraries. Coils were combined via signal-to-noise ratio (SNR) weighting, with weights computed from the reference water and noise scans, using a singular value decomposition algorithm. Spectra were aligned and phase-corrected relative to each other using a previously published robust iterative algorithm^34^. Global zero-order phase-correction was carried out based on the 3.0 ppm creatine peak in the summed spectra. No apodization or zero filling were employed. SPM12 (Wellcome Center for Human Neuroimaging, UCL, UK, http://www.fil.ion.ucl.ac.uk/spm) was used to segment the T1-weighted anatomical images into grey matter (GM), white matter (WM), and cerebrospinal fluid (CSF) images. Tissue fractions within the spectroscopic voxel were computed using VDI for subsequent use in absolute quantification and as a quality assurance metric. Metabolite quantification was carried out using LCModel^35^ version 6.3c, with a basis set containing 17 metabolites: aspartate (Asp), ascorbic acid (Asc), glycerophosphocholine (GPC), phosphocholine (PCh), creatine (Cr), phosphocreatine (PCr), GABA, glucose (Glc), glutamine (Gln), glutamate (Glu), myo-inositol (mI), lactate (Lac), N-acetylaspartate (NAA), N-acetylaspartylglutamate (NAAG), scyllo-inositol (Scyllo), glutathione (GSH), and taurine (Tau). Basis functions were simulated by solving the quantum mechanical Liouville equation using VDI, taking into account the full 3D spin profile and the actual pulse waveforms. Absolute quantification was carried out by correcting the metabolite concentrations provided by LCModel for tissue fractions estimated from the segmented images ^36^, assuming a water concentration of 43.3 M in GM, 35.88 M in white matter (WM) and 5.556 M in cerebrospinal fluid (CSF). Relaxation correction assumed the same value of T2 for GM and WM. We also assumed no metabolites in CSF tissue fractions^37^. The long TR eliminated saturation effects and, consequently, no T1 corrections were required. In addition to the concentration, the relative Cramer Rao Lower Bound (%CRLB) for each metabolite was also obtained.

#### 2.4.3 fMRI analysis

##### 2.4.3.1 Pre-processing

Functional MRI pre-processing was carried out using the FEAT tool in FSL 6.05 (FMRIB’s Software Library, www.fmrib.ox.ac.uk/fsl). The first 5 TRs of the functional data were discarded to allow steady-state magnetization. Registration of the functional data to the high resolution structural images was carried out using boundary based registration algorithm^38^. Registration of high resolution structural to standard space (1 mm MNI152) was carried out using FLIRT^39,40^, and then further refined using FNIRT nonlinear registration. Motion correction of functional data was carried out using MCFLIRT^40^, non-brain tissue removal using BET^41^, grand-mean intensity normalisation of the entire 4D dataset by a single multiplicative factor, and high-pass temporal filtering was performed with a Gaussian-weighted least-squares straight line fitting with a cut-off period of 100 seconds. Since we utilized a multivoxel-based pattern analysis (detailed below), minimal spatial smoothing using a Gaussian kernel of 2 mm FWHM was applied to the data ^42^. EPI distortion correction of the functional data was carried out with FSL-TOPUP using the acquired spin-echo field maps. In addition, independent component analysis (ICA)-based exploratory data analysis was carried out using FSL’s Multivariate Exploratory Linear Decomposition into Independent Components (MELODIC)^43^, in order to investigate the possible presence of unexpected artefacts or activations. Then, the ICA with automatic removal of motion artifacts (ICA-AROMA) tool ^44,45^ was implemented on the subject-specific spatial ICs and associated time-courses to identify motion-related noise components. The ICA-AROMA denoising strategy identifies ICA noise components based on their location at brain edges and CSF, high frequency content, and correlation with realignment parameters resulting from initial motion correction. ICA-AROMA procedure resulted in a denoised 4D time-course for each participant that was further used in subsequent task and resting-state analyses.

##### 2.4.3.2 Task-based analysis

First-level analysis was carried out on the pre-processed data for each participant. Time-series statistical analysis (pre-whitening) was carried out using FILM with local autocorrelation correction ^46^. First-level task regressors were defined based on the onset times of the task blocks and were convolved with a double-gamma hemodynamic response function. Specifically, in the current experiment we have defined two task regressors of interesting representing the early-phase and late-phase learning stages in the first day. Regressor of the temporal derivative of the task timing was also included in the analysis. Subject-specific Z-maps were thresholded non-parametrically using clusters determined by Z>3.1 and a corrected cluster significance threshold of *p*=.05. Group-level analysis was carried out using FLAME^47–49^ (FMRIB’s Local Analysis of Mixed Effects) stage 1. Group-level Z-maps were thresholded non-parametrically using clusters determined by Z>3.1 and a corrected cluster significance threshold of *p*=.05^50^. Multiple comparisons correction was spatially restricted to include only grey matter voxels using a MNI152 grey matter mask.

##### 2.4.3.3 Regions-of-interest definition

Three functional regions of interest were defined in the current study for the evaluation of M1’s functional processing characteristics, i.e., M1 and the right/left putamen. M1 ROI was initially defined as the caudal part of the precentral gyrus (the anterior wall of the central sulcus), which directly controls finger movements ^51^. This part of the motor cortex was previously shown to be functionally and anatomically differentiated from the more anterior part of the precentral gyrus corresponding to the adjacent dorsal premotor cortex ^51–53^. Then, as the study aim was to investigate aspects of a motor memory formation in M1, we defined the final M1 ROI to be the set of voxels within the caudal part of the precentral gyrus that demonstrated statistically significant increased activation during both the last (12^th^) practice block of the first day and the testing block on the second day. This was based on the definition of memory engram cells ^54,55^, i.e., activated by a learning experience, and reactivated by subsequent memory reactivation/retrieval, and also previously demonstrated in M1 following motor learning ^7^. Importantly, the M1 ROI voxels were located within the MRS voxel. Right and left ROIs of the Putamen was defined in the same way **(Figures 4A & 5A)**. These ROIs were then used in the subsequent functional analyses.

##### 2.4.3.4 Functional processing of M1

Following pre-processing, the standard space denoised time-courses were used to measure the intra- and inter-regional functional connectivity of M1 using *CONN Toolbox* v.21a (http://nitrc.org/projects/conn). Resting-state data were further denoised by regressing out the signal of the first component of the CSF ^44^ extracted with the component-based method (CompCor) implemented in CONN.

###### 2.4.3.4.1 Multivoxel local correlation (MVLC) analysis

We used the Integrated Local Correlation (ILC) analysis implemented in CONN to construct the MVLC pattern of the voxels within the M1 ROI. The ILC analysis yields the local connectivity of each voxel with its surroundings, which is characterized by the strength and sign of correlation between a given voxel’s time-course and the neighbouring voxels’ time-courses ^56–58^. We used a 1 mm kernel for characterizing the size of the local neighbourhoods, in order to express the local correlation of each voxel with its directly surrounding voxels (as the standard space data resolution was 1 mm^3^). Each voxel’s ILC value (i.e., correlation value) was then Fisher Z-transformed. Task-related MVLC pattern was defined as the spatial pattern of ILC values across all M1 ROI voxels during the late phase of learning on the first day. This is based on recent findings suggesting that activity patterns at reactivation correspond to those at the late (but not the early) phase of motor learning in M1^7^. Resting MVLC patterns were defined as the spatial pattern of ILC values across all M1 ROI voxels during either the pre- or post-learning rest periods (in which resting-state BOLD fMRI data were acquired). Next, we measured the similarity between the task MVLC pattern and the pre- and post-learning resting MLVC patterns. This was carried out by transforming the MVLC patterns to vectors and calculating the Pearson correlation between them, i.e., post rest-task correlation and pre rest-task correlation (note that the same elements across vectors correspond to the same M1 ROI voxels). Then, we computed the difference between the two similarity measures (i.e., subtracting the post rest-task similarity and the pre rest-task similarity), resulting in a similarity difference measure for each participant. This in turn enabled us to examine whether post-learning resting MVLC patterns were more similar to the task MVLC patterns compared to the pre-learning rest, therefore presumably reflecting offline memory reactivation^16^) (**Figure 4A**).

###### 2.4.3.4.2 M1-putamen functional connectivity analysis

The functional connectivity between M1 ROI and the putamen’s ROIs was calculated using an ROI-to-ROI approach, correlating the average time-courses of the ROIs at rest both before and after the task on the first day, and before the task on the second day. Each correlation value was then Fisher Z-transformed. Differences in functional connectivity were computed by subtracting post and pre values for each participant.

### 2.5 Structural MRI analysis

Overnight changes in M1 grey matter volume were examined using the voxel-based morphometry pipeline implemented in FSL (https://fsl.fmrib.ox.ac.uk/fsl/fslwiki/FSLVBM)^59,60^. FSL-VBM pre-processing first included non-brain tissue removal using BET ^41^, and tissue-type segmentation via a the Automated Segmentation Tool (FAST) to segment the images into GM, WM, and CSF. FSL FAST also performed bias-field correction for RF/B1-inhomogeneity. Next, the segmented native-space GM images were non-linearly registered with FNIRT to the standard MNI space using the ICBM-152 GM template, in order to create a study-specific GM template. Then, the GM images were non-linearly registered to the study-specific template using FNIRT. Finally, the resulting GM images were modulated by multiplying each voxel in each GM image by the Jacobian of the warp field in order to compensate for the contraction/enlargement due to the non-linear component of the transformation ^61^. M1 GM values were extracted from these non-smoothed modulated images by computing the averaged value across all voxels in the M1 ROI.

### 2.6 Statistical analyses

Statistical analyses and visualizations were performed and constructed with MATLAB, and R 4.1.2. Here, we evaluated post-learning changes (e.g., changes in metabolites concentrations, functional connectivity, behavioural performance) with linear-mixed models using the lme4 package implemented in R^62^. Each mixed effect model in the current study was examined as random intercept and random slope model, enabling the expression of different baseline levels but also difference in the extent of change for each evaluated measure across the participants. To this end, participants ID was used as the random effect and time as the fixed effect: *dependent variable* ∼ *time* + (*Time* | *ID*). Pairwise comparisons in which statistically significant effects were observed were corrected for multiple tests using the False Discovery Rate (FDR) ^63,64^. For the MRS data, the mean concentration and standard deviation of each metabolite, as well as the mean and standard deviation of the CRLB were calculated for each of the time points. Quality assurance of the MRS data followed previously reported metrics: visual inspection for gross artifacts, such as lipid contamination and spurious echoes, metabolites concentrations that were three standard deviations away from the mean of all time-points measurements were excluded from further analyses, as well as water linewidths exceeding 15 Hz FWHM, and SNR of ≤30 from the LCModel output ^11,19,65^. MRS data quality summary and LCModel fitting representation are presented in the **Supplementary Table 1** and **Supplementary Figure 1**, respectively. To evaluate consistency in the MRS voxel placement across the two scanning sessions, repeated measure mixed models were used for GM, WM, and CSF tissue fractions of the two voxels. These comparisons did not demonstrate statistically significant tissue fraction differences between the scanning sessions (*See Supplementary Material*). Relationships between continuous variables were assessed using Pearson’s correlation coefficient (two-tailed tests). Since we aimed to investigate the unique relationship between Glu or GABA and the other brain/behaviour changes, we computed partial correlations while controlling for the other metabolite. The associations between the proposed consolidation and plasticity measures (i.e., functional connectivity metrics and GM volume) and the change in behavioral performance overnight were evaluated using a one-tailed correlation test as directional relationships were hypothesized as part of the rationale of the study. All correlations evaluating a relationship with overnight performance changes were adjusted for performance on the last practice block on the first, as higher performance on the first day was associated with smaller overnight gains in performance (*p*<.05).

## 3. Results

### 3.1 Increased coupling between glutamate and GABA following learning

Glu levels were found to exhibit a general trend of increase during the period following learning (*F*(5,33.8)=2.14, *p*=.084) (**Figure 2A**). Pairwise comparisons between each post-learning measure and pre-learning levels demonstrated a statistical trend for increased Glu only after 30 minutes following learning (β=.27, *p*=.074 FDR-corrected, *p*=.014 uncorrected), which corresponded to an increase of ∼3.4%. GABA levels, on the other hand, were not found to be statistically different on each time point following learning compared to baseline levels (*F*(5,34.7)=0.82, *p*=.544) (**Figure 2A**). Also, while Glu showed a general pattern of monotonic increase during this period, mean GABA levels fluctuated with no clear trend. Resting levels on the second day were also not statistically different from pre-learning baseline levels on day 1 for both metabolites (**Figure 2A**).

**Figure 2.**
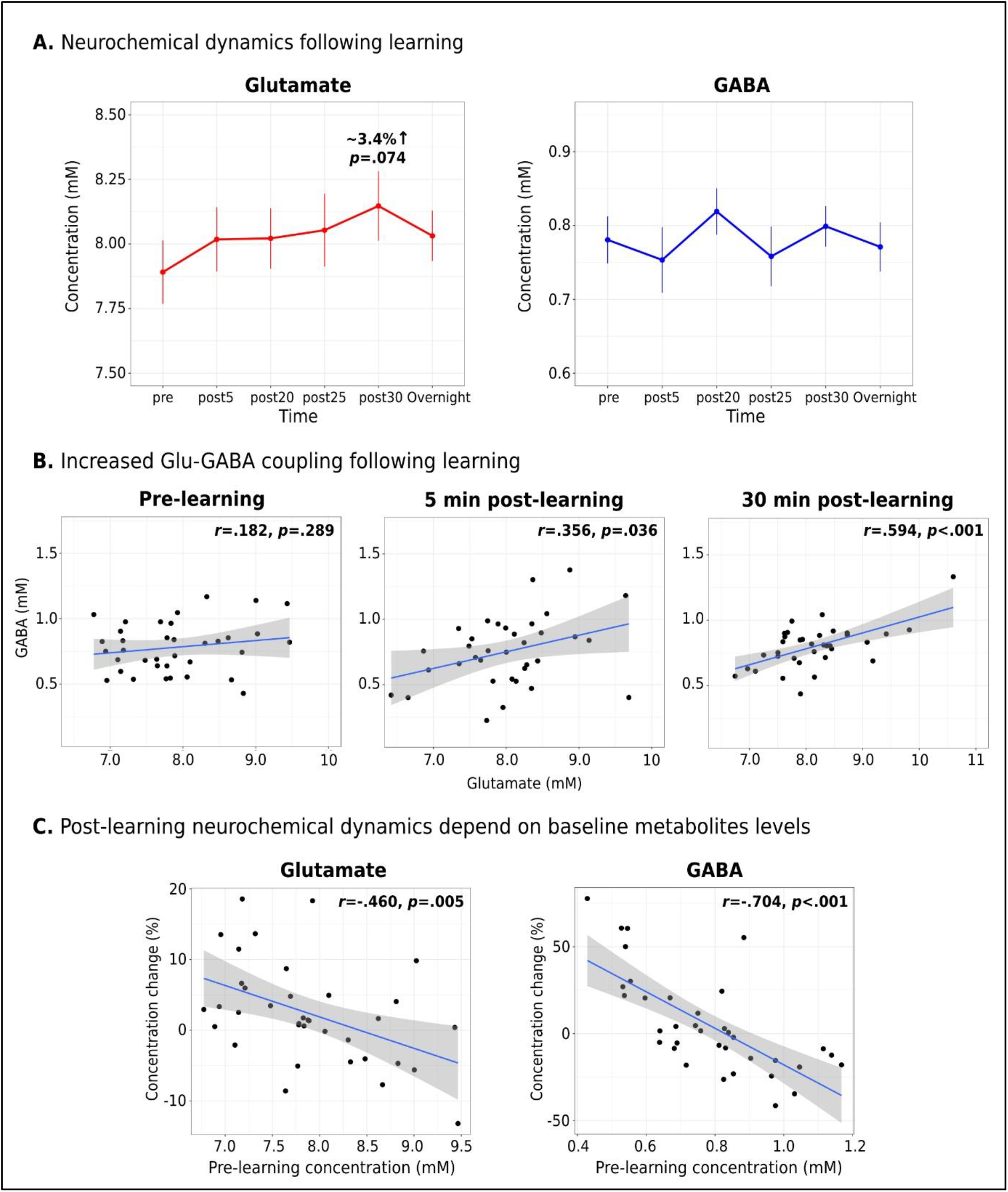
Neurochemical dynamics following motor learning as reflected by changes in Glu and GABA concentrations during consolidation and overnight (**A**); Dynamics of the “balance” between excitation and inhibition in M1 across participants following learning (**B**); Relationship between baseline metabolite concentrations and the extent of concentration change following learning (**C**).

**Figure 3.**
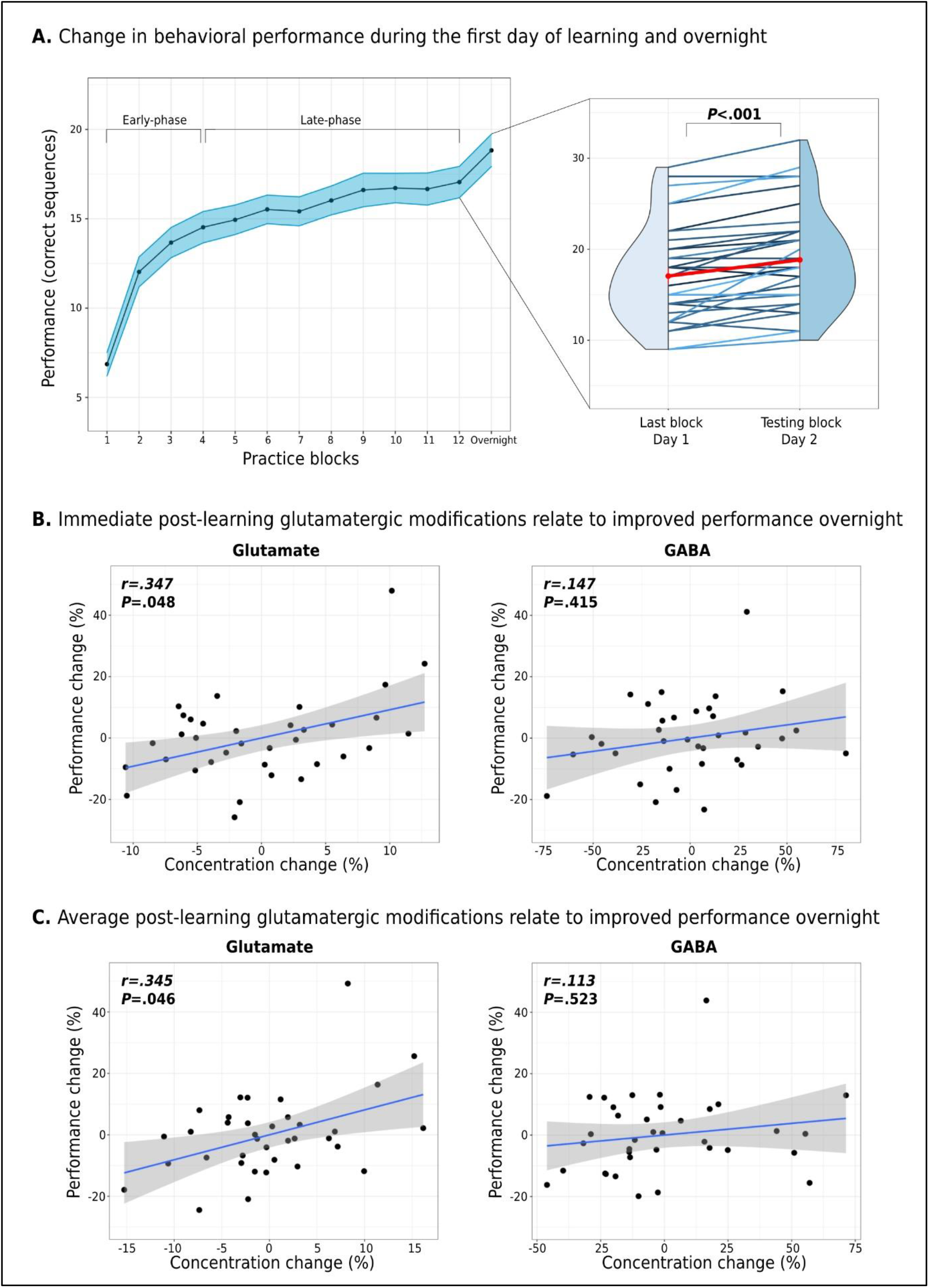
Neurochemical dynamics and behavioral performance. Behavioral changes in task performance during the first day of learning and overnight (**A**); The relationship between Glu and GABA changes immediately following learning and overnight changes in task performance (**B**); The relationship between Glu and GABA changes averaged across the 30-min following learning and overnight changes in task performance (**C**).

Next, we measured the dynamics of the correlation between Glu and GABA levels following learning compared to pre-learning levels, as a reflection of the balance between excitation and inhibition across participants prior-to and following learning. While pre-learning levels of Glu and GABA demonstrated a low, non-significant correlation (*r*=.182, *p*=.289), the relationship between the two metabolites increased during the post-learning period from (*r*=.356, *p*=.036) immediately following the task in the first 5 minutes of consolidation, to (*r*=.594, *p*<.001) after 30 minutes (**Figure 2B**). However, it should be noted that when we directly compared the pre- and post-learning correlation coefficients, only the correlation value at 30 minutes following learning was significantly higher than the pre-learning value (Fisher’s Z = 1.983, p = .024).

Lastly, we found an inverse relationship between pre-learning Glu and GABA concentrations and their dynamics following learning. Specifically, higher Glu and GABA at baseline were both associated with a greater decrease in metabolites levels during the 30 minutes following learning, and vice versa (*r* = -.460, *p* = .005, and *r* = -.704, *p*<.001, for Glu and GABA, respectively) (**Figure 2c**).

### 3.2 Increased glutamate following learning support behavioral improvement overnight

A significant overnight improvement in skill performance of 11.82±14.3% was found at the group-level between the last practice block on day 1 and the testing block on day 2 (*F*(1,35)=33.12, *p*<.0001), although some participants did demonstrate declined or stable performance between the two sessions (**Figure 3A**). When examining the relationship between post-learning neurochemical modifications and overnight changes in performance, only increased Glu, both immediately after learning and averaged across the 30-min post-learning, was related to greater overnight behavioral improvements (*r*=.347, *p*=.048 and *r*=.345, *p*=.046, respectively) (**Figure 3B-C**).

**Figure 4.**
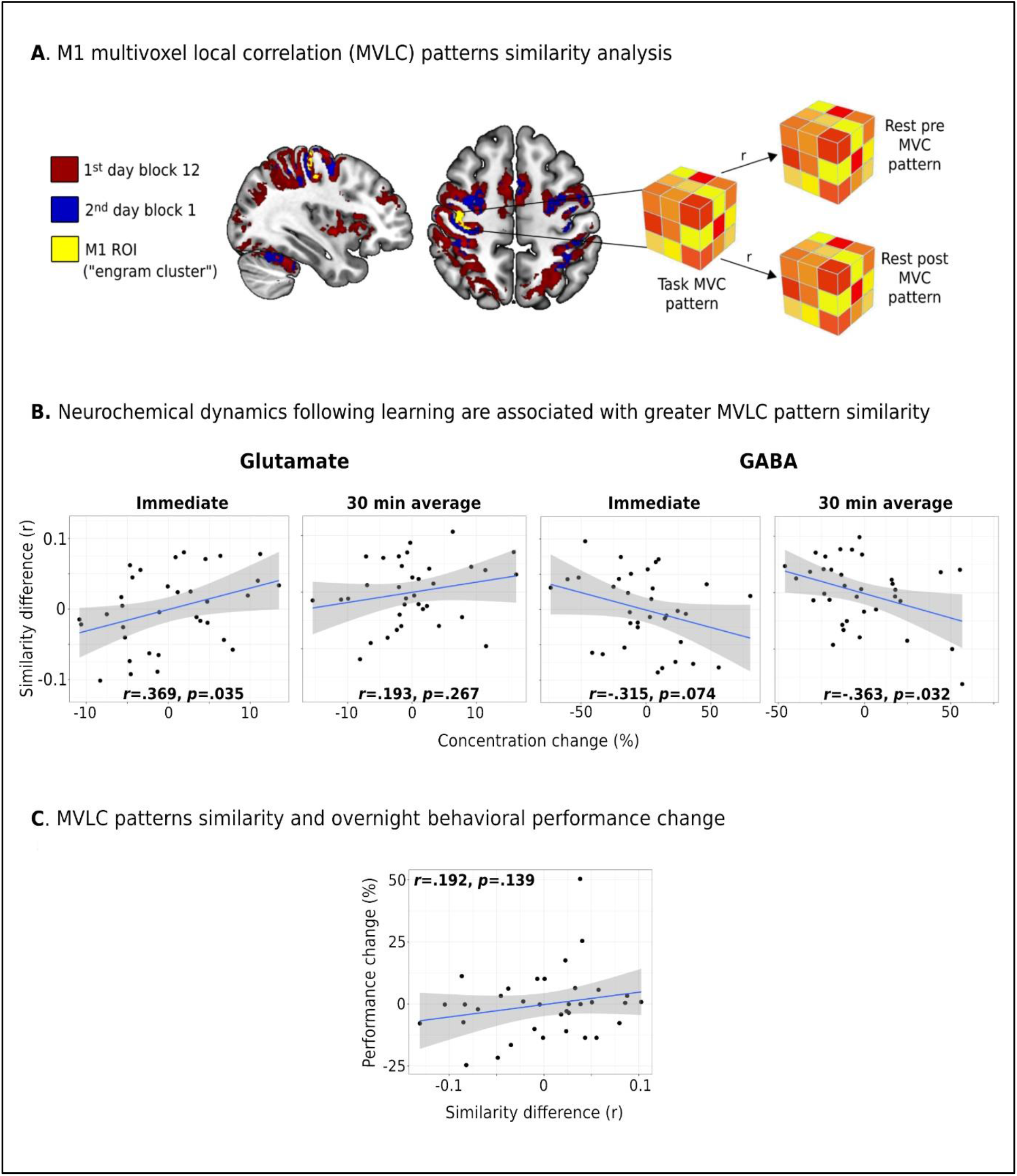
Multivoxel local correlation patterns and neurochemical dynamics. MVLC analysis and M1 ROI definition (yellow) based on activation patterns on day 1 (red) and 2 (blue) (**A**); The relationship between MVLC similarity differences and Glu and GABA changes post-learning (**B**); The relationship between MVLC similarity difference and overnight change in task performance (**C**).

**Figure 5.**
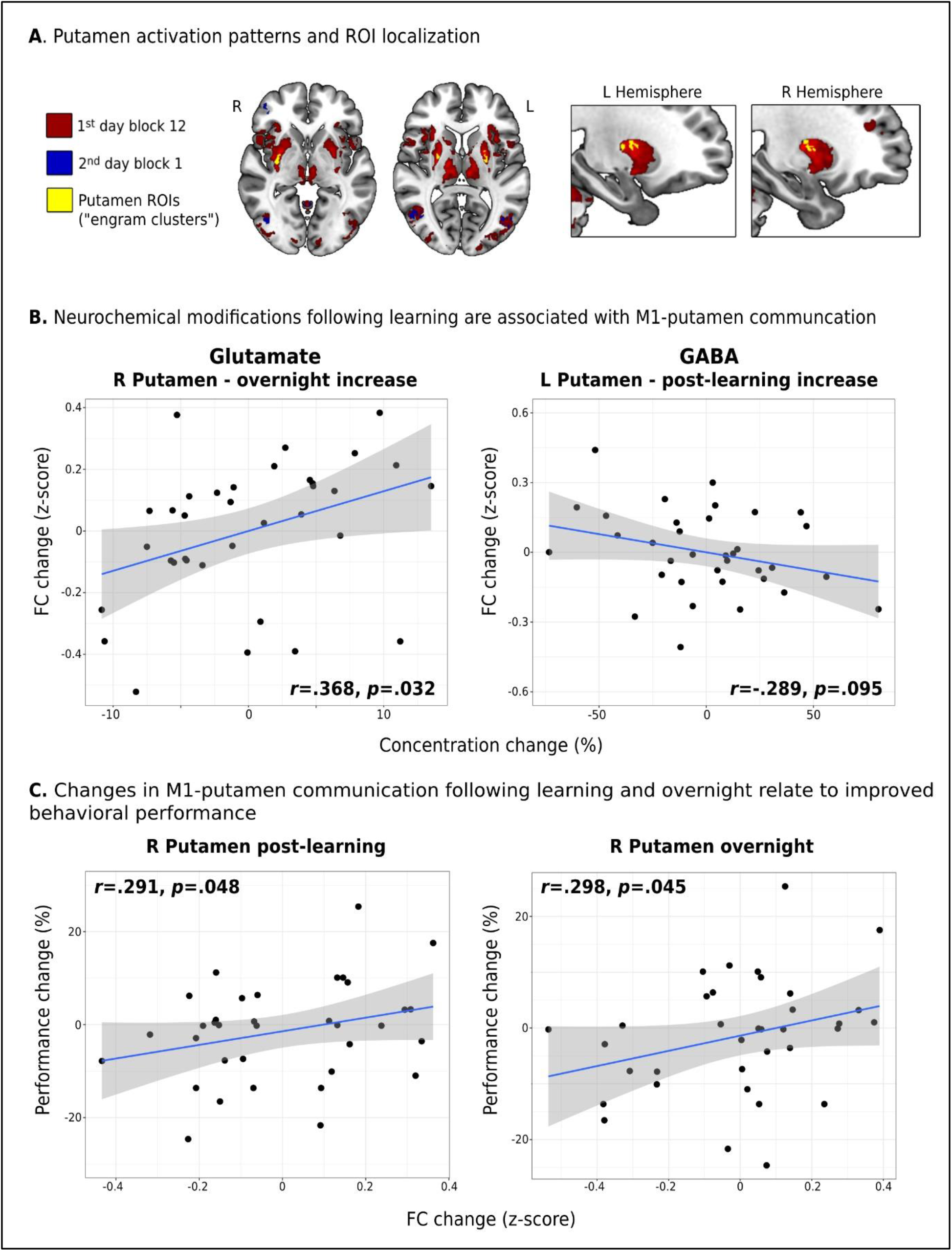
Neurochemical dynamics following learning and M1 functional connectivity with the putamen. Putamen ROI definition (yellow) based on activation patterns on day 1 (red) and 2 (blue) (**A**); The relationship between immediate post-learning changes in Glu and GABA changes and changes in M1-Putamen functional connectivity (**B**); The relationship between M1-Putamen functional connectivity and overnight changes in task performance (**C**). FC = functional connectivity; L = left; R = right

### 3.3 Increased excitation relates to greater MVLC pattern similarity following learning

Group level repeated measure mixed model analysis did not reveal a significant difference between the similarity of the post-learning rest MVLC patterns with the task MVLC pattern compared to the pre-learning rest (*F*(1,35)=0.03, *p*=.876). Thus, at the group level we did not find evidence for offline motor memory reactivation based on the MVLC patterns. However, higher similarity of the post-learning rest with the task compared to pre-learning patterns was associated with greater increases in Glu immediately following learning (*r*=.369, *p*=.035), and with greater reduction in GABA both immediately after learning (*r*=-.315, *p*=.074) and averaged across the 30-min following the task (*r*=-.363, *p*=.032) (**Figure 4B**). In addition, we did not observe a significant relationship between the MVLC similarity difference and the change in performance overnight (*r*=.192, *p*=.139) (**figure 4C**).

### 3.4 Increased glutamate following learning supports increased M1-putamen functional communication

We found a trend towards an association between increased M1 connectivity with the left putamen and reduced GABA immediately following learning and (*r*=-.289, *p*=.095, **Figure 5B**), but not the right putamen (*r*=-.197, *p*=.265). Immediate changes in Glu were not related to changes in M1-putamen connectivity (right putamen: *r*=.002, *p*=.992; left putamen: *r*=-.031, *p*=.862). Average changes in Glu (right putamen: *r*=.054, *p*=.760; left putamen: *r*=.052, *p*=.768) or GABA (right putamen: *r*=-.035, *p*=.841; left putamen: *r*=-.229, *p*=.186) across the 30 minutes after learning were also not associated with connectivity changes with the putamen. While a group-level analysis did not reveal significant differences in functional connectivity of M1 with either the right or left Putman when comparing the post-learning resting period with pre-learning rest (right Putamen: *F*(2,35)=1.51, *p*=.141, left Putamen: *F*(2,35)=1.39, *p*=.247), greater increases in the functional connectivity of M1 with the right putamen were associated with greater overnight improvements in skill performance (*r*=.291, *p*=.048, **Figure 5C**), a relationship which was not observed with the left putamen (*r*=-.029, *p*=.870).

### 3.5 Neurochemical modifications following learning support overnight functional and structural plasticity in M1

#### 3.5.1 M1 functional connectivity with the putamen

When examining the relationship between post-learning neurochemical changes and overnight changes in the functional connectivity of M1 with the putamen, we found increased connectivity with the right putamen to associate with increased Glu immediately following learning (*r*=.368, *p*=.032), but not with GABA (*r*=-.141, *p*=.426) (**Figure 5B**). Changes in Glu or GABA immediately after learning were not significantly associated with overnight changes in connectivity with the left putamen (Glu: *r*=.130, *p*=.465; GABA: *r*=-.074, *p*=.677), and there were also no significant correlations between average post-learning Glu or GABA changes with overnight changes in the connectivity with either the right (Glu: *r*=.259, *p*=.132; GABA: *r*=.185, *p*=.287) or the left putamen (Glu: *r*=.144, *p*=.410; GABA: *r*=.250, *p*=.147) (**Supplementary Figure 2**). Furthermore, while overnight connectivity values were not significantly different from pre-learning values at the group-level (right putamen: *F*(2,35)=0.73, *p*=.470; left putamen: *F*(2,35)=0.35, *p*=.727), overnight increases in the connectivity with the right putamen (*r*=.298, *p*=.045), but not the left putamen (*r*=.156, *p*=.372), were significantly associated with greater improvements in task performance (**Figure 5C**).

#### 3.5.2 M1 grey matter volume

When we examined the relationship between post-learning neurochemical modifications and overnight changes in M1 GM volume we found reductions in GABA concentrations during the 30-min window post-learning were associated with overnight increases to M1 GM volume (*r*=-.349, *p*=.043) (**Figure 6A**). Although the average post-learning change in Glu was also negatively correlated with increased M1 GM volume, it was not statistically significant (*r*=-.202, *p*=.252). Since increased M1 GM volume was correlated with decreased levels of both GABA and Glu (although to different extent) we also examined the relationship between changes in M1 GM volume and the ratio of Glu and GABA (Glu/GABA) as a correlate of excitation-inhibition balance. This analysis revealed a significant correlation between increased Glu/GABA ratio following learning (i.e., greater excitation/disinhibition) and overnight increases in M1 GM volume (*r*=.340, *p*=.049) (**Figure 6A**). Changes in Glu or GABA immediately following learning were not significantly related to overnight changes in M1 GM volume (Glu: *r*=-.182, *p*=.310; GABA: *r*=-.012, *p*=.949). While we did find a link between overnight changes in M1 GM volume and post-learning neurochemical modifications, there was not a group-level change in M1 GM volume overnight (*F*(1,35)=1.59, *p*=.215)), and M1 structural changes were not associated with overnight changes in task performance (*r*=-.124, *p*=.242) (**Figure 6B**).

**Figure 6.**
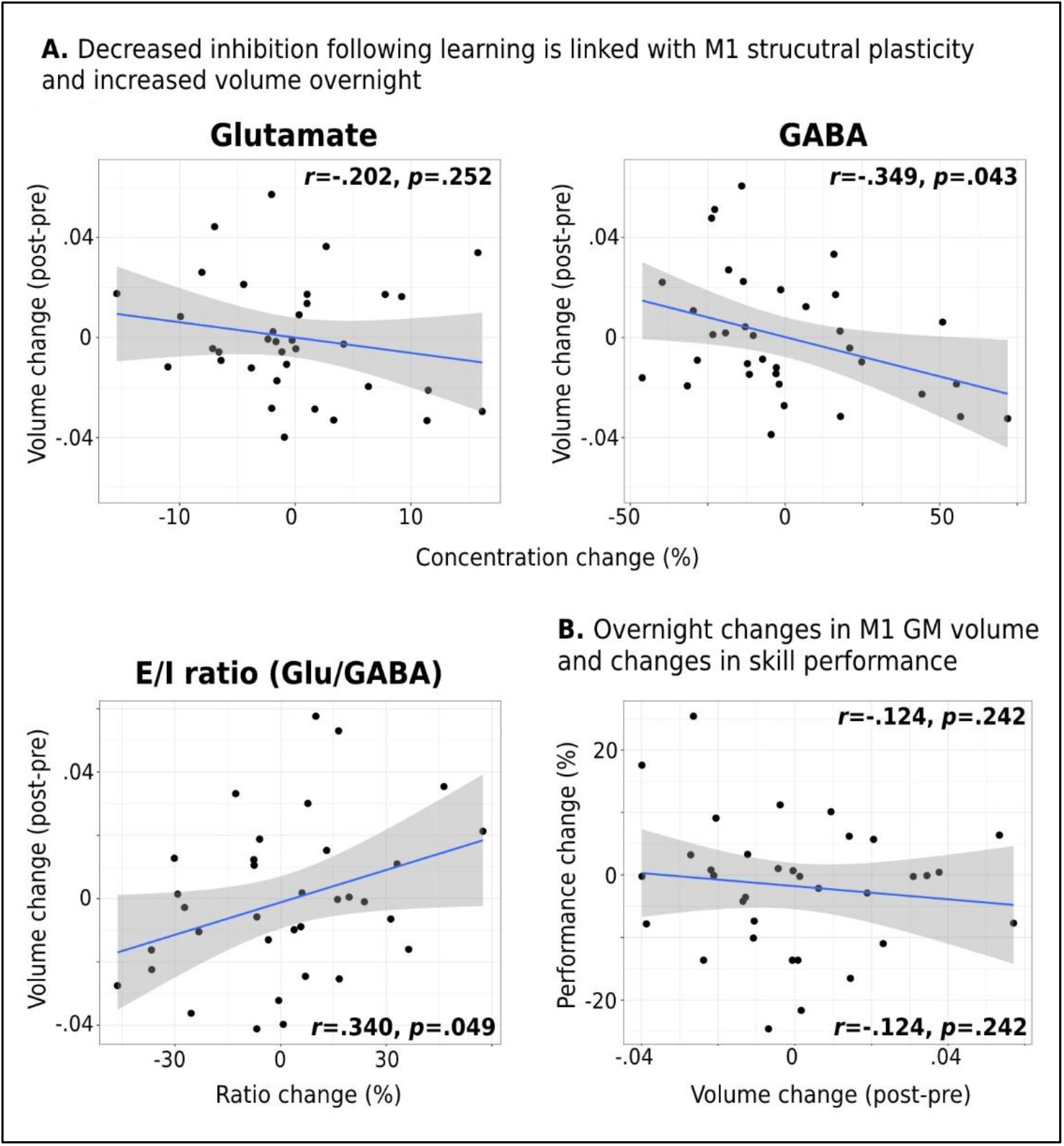
Neurochemical dynamics and M1 structural plasticity. The relationship between Glu, GABA, and E/I ratio post-learning changes and changes in M1 GM volume overnight (**A**); The relationship between overnight M1 GM volume changes and changes in task performance (**B**).

## 4. Discussion

### 4.1 Glu-GABA Coupling in M1 Increases Following Motor Learning

The aim of current study was to reveal the dynamics of Glu and GABA in M1 during the early stages of motor memory consolidation, and how these may relate to skill learning-induced neuro-behavioral plasticity. We also examined how early changes in Glu and GABA following learning may relate to local and inter-regional functional processing of M1. These questions follow previous findings suggesting that the initial first hours after skill learning may be vital for the consolidation of motor skills^5,9,27,66^. Interestingly, at the group-level, we did not find significant changes in either Glu or GABA, either immediately or during the first 30 minutes following the task, despite observing a trend for increased Glu after 30 minutes. This is partially in contrast to a recent study by Maruyama and colleagues^11^ which found decreased Glu immediately after learning. However, the reduction in Glu observed in their study was similar in both the learning group and a non-learning control group, suggesting that it may not have represented a learning-induced physiological process. Nonetheless, while not observing group-level changes in Glu or GABA dynamics, we did reveal increased coupling between these metabolites during the first 30 minutes of offline consolidation following learning. The weak correlation between Glu and GABA levels observed at baseline prior to the learning task is in accordance with previous findings in the motor cortex^67^, raising the possibility that the observed increase in Glu-GABA coupling after the task may be a consequence of the learning experience. Previous works had demonstrated changes in E-I during the online phase of motor learning, mainly expressed as decreased GABA levels^19,24^. Although we did not measure changes in Glu and GABA during the learning phase, it is possible that the increase in Glu-GABA coupling following the task may represent a physiological “rebound” effect aiming to restore baseline physiological functions after being perturbed during the learning itself. On the other hand, as the E-I balance has been proposed to modulate important aspects of cognition and behavior^17^, the increased Glu-GABA coupling may reflect a direct learning-related phenomenon, presumably reflecting a more controlled or fine-tuned processing of the learned experience in the motor cortex during the early stages of consolidation. It was recently suggested that the E-I balance may be vital for neuronal computation that is robust to noise and for shaping efficient neural coding^68,69^. Furthermore, E-I balanced neural circuitry have been linked with the storage of short-term memory and the stability of memory states^68,70^. As motor memories are significantly vulnerable to disruption and interference during the initial offline period immediately following learning, increasing the Glu-GABA coupling may subserve the initial stages of motor consolidation that are critical for the stabilization of the motor memory trace^5,6^.

### 4.2 Early Post-Learning Glutamate Changes Support Overnight Motor Skill Consolidation

While we did not observe a mean change in neurochemical dynamics following learning across the whole sample, we revealed an association on a subject-by-subject basis between increased Glu (both immediately and 30 minutes following learning) and greater overnight improvements in skill performance. It is well established that glutamatergic neurons are strongly implicated in the induction of LTP, one of the most-studied forms of synaptic plasticity^71^. LTP-like plasticity has been shown to be induced in the rodent motor cortex artificially and following motor learning^20,72,73^, and was also demonstrated in the human M1 following motor learning using non-invasive brain stimulation^74^. While LTP-dependent synaptic strengthening has mostly been associated with post-synaptic modifications in Glu receptors, LTP-dependent increased synaptic efficacy may also be expressed by an increase in the release of Glu by the pre-synaptic neuron, and therefore an increase in the amount of the transmitter outside the vesicles^75–77^. Increased Glu release was also found to occur in the rodent motor cortex following motor learning^20^. It has been proposed that the vesicular pool is largely invisible to MRS compared to the extracellular and cytosolic pools of neurons and astrocytes. Therefore, a presynaptic LTP-related shift of more Glu into MR-visible compartments (i.e., extra-vesicular) may underlie the behaviorally beneficial increased MRS Glu signal observed following learning, similar to what have been proposed to occur during task-related functional activation^78^. This hypothesis, however, may need further examination, preferably using animal models. In addition to LTP, increased cortical excitability following motor learning has been linked to offline behavioural improvements. Since increased excitability may result from either increased Glu or decreased GABA, the behaviorally beneficial post-learning increases in Glu exhibited in the current study may therefore represent a form of increased excitability in M1. Interestingly, while we have not seen significant changes in M1 neurochemistry (as well as in function or structure) at the whole sample level, we did find significant group-level increases in skill performance overnight. This discrepancy may result from the fact that the current study focused on potential consolidation mechanisms that are expressed during wakefulness, and not during sleep, which has been shown to play a key role in the performance-enhancing effect of consolidation following explicit motor learning (as been implemented in the current work) ^5,79^. Physiological phenomena taking place during sleep such as sleep spindles have been implicated in plastic processes and memory enhancement and may be responsible (at least in part) for the increased performance that was demonstrate for most of the participants, despite not exhibiting significant changes in neural mechanisms of consolidation during wakefulness.

### 4.3 Increased Cortical Excitation Is Linked with M1 Functional Processing and Plasticity Following Learning

Increased cortical excitability has been proposed to affect neural processes that may be essential for the development of offline behavioral improvements and cortical plasticity. One of these processes may be the offline reactivation/replay of recently acquired motor memories. Offline memory reactivation/replay is currently thought to play a vital role in the consolidation of explicit and implicit new memories^14^, even during wakefulness^80^. While offline memory reactivation has been investigated mainly in the hippocampus and in the context of episodic memory, recent evidence suggest a similar process occurs in the motor cortex in both animals^15^ and humans^3^. Here, we found that both increased Glu and decreased GABA were independently associated with higher similarity of M1 MVLC resting patterns immediately following learning to the patterns observed during the learning experience itself, potentially reflecting a process of offline motor memory reactivation. Intra-regional patterns of multivoxel functional connectivity were previously used to examine offline memory reactivation in humans^16^, although these studies have focused on different brain regions^28^ (e.g., the hippocampus). Furthermore, although the current study did not utilize a direct measure of neuronal reactivation/replay, the recent evidence for a motor memory-related replay in M1 at the cellular level following motor learning in rodents ^15^ may support the feasibility of this process in the human motor cortex as well.

In addition to the role of glutamatergic modifications in synaptic plasticity, GABAergic modulation has also been implicated in motor learning and plasticity processes. Evidence from the animal literature have demonstrated GABAergic disinhibition to be essential for the induction of LTP-like plasticity and the facilitation of LTP-like synaptic responses in M1 ^21,81^. Animal studies have also shown motor cortical disinhibition at the cellular level during and following motor practice that is expressed by rapid elimination of inhibitory boutons on layer II/III excitatory neurons ^82^ and decreased release of GABA ^20^. Studies in humans utilizing MRS have demonstrated a reduction in cortical GABA during motor learning, but not during plane movements ^19,24^, suggesting a learning-specific modulation of GABAergic function. The relationship observed in the current study between GABAergic reduction and our MVLC correlate of memory reactivation may provide important insights into the potential role of GABAergic modulation in offline motor learning processes, in addition to its role during the learning experience itself.

Previous studies in humans have also demonstrated a potential link between local GABA levels and functional connectivity of the motor cortex. Specifically, lower GABA concentrations were found to associate with increased resting state functional connectivity of the motor network^83^, while decreased GABA-to-Glu ratio (i.e., disinhibition) was associated with greater increases in M1 functional connectivity with the frontoparietal network following motor learning^11^. Here, we found a trend towards a relationship between reduction in GABA immediately following learning and increased functional connectivity between M1 and the contralateral putamen. We also found a relationship between immediate Glu increase after learning and overnight increase in M1 connectivity with the right putamen; However, we did not observe a significant group-level increase in M1-putamen connectivity, neither during the 30 minutes of consolidation or overnight. Motor learning has been shown to induce strengthening of M1 engram neurons synaptic outputs to the striatum, which may be modulated by changes in the neuronal output of M1 excitatory cells at striatal dendrites ^7^. These modifications in turn would have been neurochemically expressed remotely of the motor cortex, and therefore cannot directly explain the relationship observed in the current study between the Glu increase at the site of M1 and the increased M1-putamen connectivity. Nevertheless, the relationship shown in the current work between M1-right putamen functional connectivity (both during consolidation and overnight) and the overnight behavioral improvements further highlights the importance of this circuit in the consolidation of new motor memories. Furthermore, it complements previous findings which highlight the importance of immediate post-learning functional connectivity modifications in M1 to motor memory consolidation ^9^.

### 4.4 Reductions in M1 GABA Precede and Support Overnight Plasticity-Related Volume Changes

In addition to learning-induced functional synaptic strengthening, synapses also store information by modifying their structure. Increased protein synthesis and dendritic spines remodelling have been well documented during the late phases of LTP, resulting in enlargement of existing spines or the formation of new ones ^71^. Furthermore, motor learning has been shown to promote the formation and stabilization of new spines in M1, that is expressed specifically in M1 engram cells ^7^. Previous studies have shown that intense local release of Glu or GABA can induce post-synaptic dendritic spines formation ^18^. By utilizing two-photon uncaging of caged glutamate compounds, which reliably stimulates single spines, it was shown that releasing Glu into the synaptic cleft and maximally activating the Glu NMDA receptors, which are critical for LTP induction ^71^, induce robust spine enlargement in the adult mouse neocortex ^84^. Interestingly, increased release of GABA was shown to induce dendritic spine enlargement only during early life development, when GABA is still a depolarizing agent, further highlighting the potential role of excitation (or disinhibition) in synaptic plasticity. These findings suggest that presynaptic neurotransmitter release may constitute the main trigger for structural synaptic plasticity and highlight the roles of Glu and GABA in promoting synaptic remodelling ^18^. Our findings follow this physiological rationale on the macroscale as we found GABAergic reduction and Glu-to-GABA ratio increase following learning to associate with overnight increases in M1 GM volume. While human structural MRI cannot compete with the microscopic resolution enabled in animal studies, cortical macrostructural changes have been demonstrated following motor learning in the human brain ^85,86^, including in the motor cortex ^8^. Yet, linking those changes to specific microscopic modifications is not trivial. Also, while the animal literature has proposed a strong link between structural plasticity and adaptive behavior following motor learning, findings in humans are less consistent in demonstrating such as relationship ^8,85^, as also apparent in the current study. This in turn may result from the limited mechanistic specificity that is inherent in structural human neuroimaging methods and makes the direct linkage between human and animal findings less straightforward. Further, learning-induced structural plasticity involve processes of formation as well as the elimination of existing synapses, of which temporal dynamics may differ across individuals, potentially also introducing inter-individual variability in the expression of macroscale plasticity. As with Glu, we did not find statistically significant group-level changes in GABA following learning, despite significant correlations between GABA dynamics and M1 structure and function on a subject-by-subject basis. GABAergic inhibitory neurons display wide variety of morphological and physiological properties. Different subtypes of GABAergic neurons target different anatomical domains of excitatory neurons, enabling them to regulate different aspects of the spatiotemporal activity of the glutamatergic cells ^87^. Therefore, GABAergic modulation following learning may be subtype-specific and function-specific, as was evidence by both increased and decreased inhibitory axonal boutons among different GABAergic cells subtypes following the same motor learning ^82^. This opposite modulatory pattern may in turn underlie a lack of absolute change in GABAergic concentration across the motor cortex, and may potentially explain why a group-level change in whole M1 GABA has not been observed with MRS following motor learning in the current and other studies ^11^. In contrast, the subject-based correlation we did find in the current study between GABA dynamics and other neural metrics may reflect the dynamics within the GABAergic pool itself across different individuals. It is also important to highlight, that while we did not observe group-level changes in Glu and GABA following learning, we did observe a continuous increase in the correlation between the metabolites’ concentrations during the post-learning period compared to the pre-learning value. This increased “coherence” between Glu and GABA levels following learning may represent greater fine-tuned regulation of the balance between inhibition and excitation that may support further consolidation processes. In addition, it could also reflect greater homeostatic control mechanism aiming to restore the balance between excitation and inhibition after it has been perturbed, for example by learning-induced disinhibition during the online learning experience ^19,24^, although both of these hypotheses require further examination.

## 5. Conclusions

Many human studies to date have focused on the ratio between Glu and GABA as a reference for the balance between excitation and inhibition. Here, we show that Glu and GABA may be independently associated with different aspects of motor memory consolidation and plasticity on different timescales, suggesting that learning-induced modifications of these metabolites may underpin distinctive functions in learning and memory processes. We have demonstrated that changes in Glu and GABA in M1 early following motor learning may be important for offline consolidation processes and the promotion of structural and functional cortical plasticity. Also, while the post-learning neurobehavioral changes were mostly associated with immediate post-learning changes in Glu, these were more linked to GABAergic changes expressed across the entire 30 minutes period after learning, suggesting that Glu- and GABA-dependent plasticity processes may operate on different timescales. Hence, the current study provides important insights to our basic understanding of the multidimensional mechanisms of learning and plasticity in the human brain. Furthermore, in addition to the derived biological insights, our findings may also have important clinical implications. Non-invasive brain stimulation methods such as transcranial magnetic stimulation (TMS) and transcranial direct current stimulation (tDCS) have been shown to be able to “externally” induce increased cortical excitability and LTP-like plasticity^88^, and to promote GABAergic and glutamatergic modulation^89^. Therefore, the potential key role of early post-learning neurochemical modifications to motor learning and plasticity that was revealed in the current study may be further examined in clinical trials and clinical settings of rehabilitation following stroke or brain injury using the methods mentioned above.

## Acknowledgment

Assaf Tal acknowledges the support of the Monroy-Marks Career Development Fund, the historic generosity of the Harold Perlman Family, and the support of NIH grant R21NS112853 and Israeli Science Foundation personal grant 416/20. Dr. E. Furman-Haran holds the Calin and Elaine Rovinescu Research Fellow Chair for Brain Research. We would like to acknowledge Edward J. Auerbach, Ph.D. and Małgorzata Marjańska, Ph.D. (Center for Magnetic Resonance Research and Department of Radiology, University of Minnesota, USA) for the development of the pulse sequences for the Siemens platform which were provided by the University of Minnesota under a C2P agreement. We also thank Prof. Nitzan Censor for his thoughtful advice and Dr. Tali Weiss for her assistance with constructing the fMRI task paradigm.

## Supplementary Material

### MRS quality metrics

The MRS quality metrics are presented in **Table S1**. Out of 216 MRS measurements across all participants (36 participants x 6 MRS runs), 5 measurements (across 3 participants) were not included in the final data analyses due to low SNR (i.e., <30) or contaminated spectrum, and 2 other measurements from an additional participant were not acquired due to time limitation in the scanner. Repeated measures mixed model analysis confirmed no significant differences in the voxels GM fraction (*p*=.601), WM fraction (*p*=.507), or CSF fraction (*p*=.560) between the two sessions. Furthermore, water linewidth was not statistically different between each pair of MRS runs over the two sessions (*p*>.05, FDR-corrected). The SNR on the second day MRS run was significantly higher compared to each post-learning run on day 1 (*p*<.05, FDR-corrected). No differences were found between each two runs within day 1 (*p*>.19 for all pairwise comparisons, FDR-corrected). **Figure S1** demonstrates a representative spectrum acquired from one participant.

**Table S1.** MRS quality metrics (Mean±SD)

| MRS run | Water FWHM | SNR | CRLB (GABA) | Voxel GM fraction | Voxel WM fraction | Voxel CSF fraction |
| --- | --- | --- | --- | --- | --- | --- |
| Pre-learning | 10.67±1.18 | 59.06±5.67 | 29.33±8.3 | 0.373±0.03 | 0.575±0.05 | 0.052±0.03 |
| Post 5 min | 10.79±1.08 | 58.83±5.68 | 33.54±15.3 | 0.373±0.03 | 0.575±0.05 | 0.052±0.03 |
| Post 20 min | 10.67±1.09 | 58.49±5.47 | 28.51±7.2 | 0.373±0.03 | 0.575±0.05 | 0.052±0.03 |
| Post 25 min | 10.81±1.22 | 57.28±6.14 | 32.90±12.6 | 0.373±0.03 | 0.575±0.05 | 0.052±0.03 |
| Post 30 min | 10.94±1.18 | 58.1±5.11 | 29.06±5.9 | 0.373±0.03 | 0.575±0.05 | 0.052±0.03 |
| Overnight | 10.58±1.09 | 60.83±5.78 | 29.86±9.7 | 0.370±0.03 | 0.579±0.05 | 0.050±0.03 |

**Figure 1.**
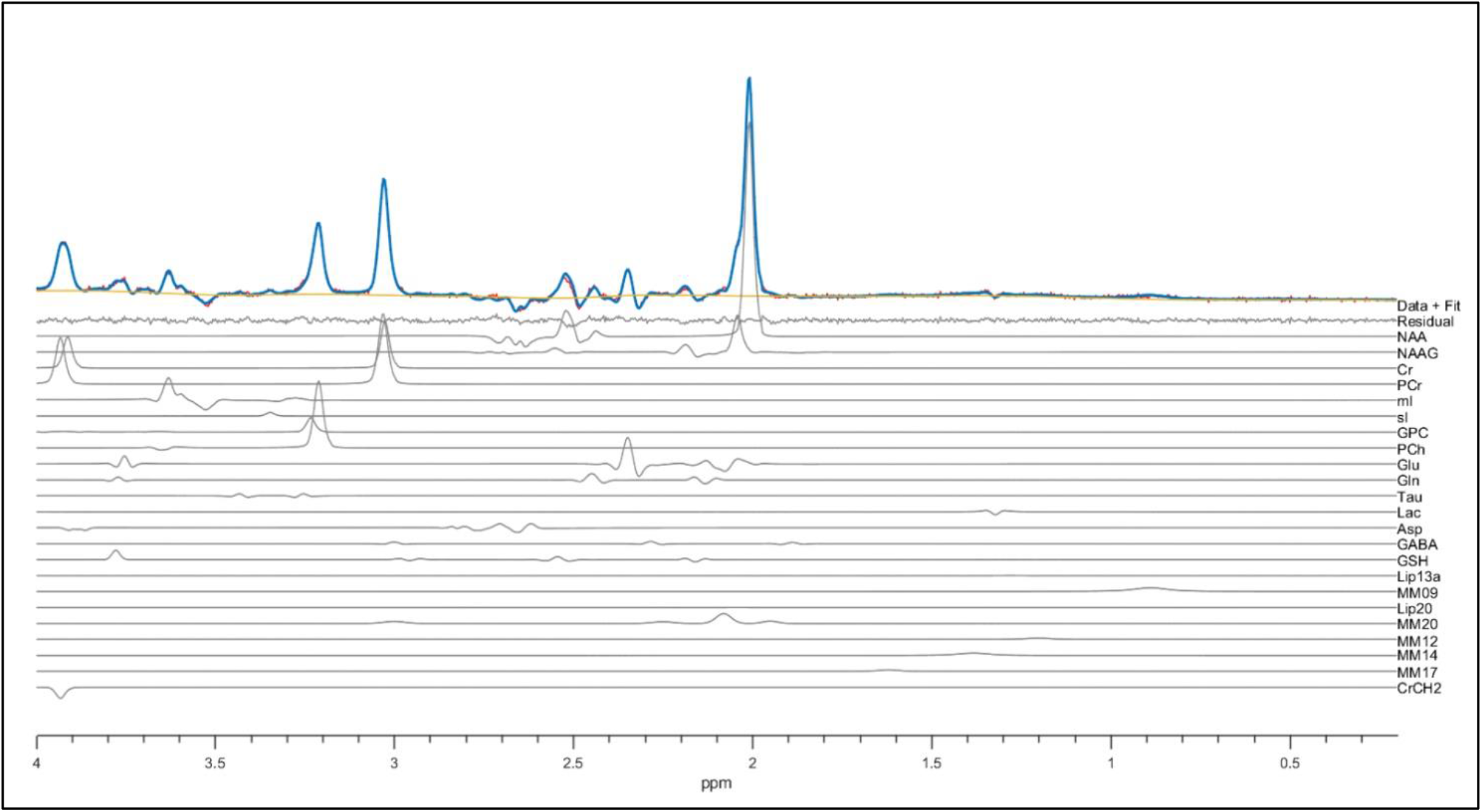
A representative spectrum acquisition from one participant including model fit.

## References

1. Dayan E, Cohen LG. Neuroplasticity subserving motor skill learning. Neuron. 2011;72(3):443–454. doi:10.1016/j.neuron.2011.10.008

2. Herszage J, Sharon H, Censor N. Reactivation-induced motor skill learning. Proc Natl Acad Sci. 2021;118(23):e2102242118. doi:10.1073/pnas.2102242118

3. Buch ER, Claudino L, Quentin R, Bönstrup M, Cohen LG. Consolidation of human skill linked to waking hippocampo-neocortical replay. Cell Rep. 2021;35(10):109193. doi:https://doi.org/10.1016/j.celrep.2021.109193

4. Breton J, Robertson EM. Flipping the switch: mechanisms that regulate memory consolidation. Trends Cogn Sci. 2014;18(12):629–634. doi:10.1016/j.tics.2014.08.005

5. Walker MP, Brakefield T, Hobson JA, Stickgold R. Dissociable stages of human memory consolidation and reconsolidation. Nature. 2003;425(6958):616–620. doi:10.1038/nature01930

6. Brashers-Krug T, Shadmehr R, Bizzi E. Consolidation in human motor memory. Nature. 1996;382(6588):252–255. doi:10.1038/382252a0

7. Hwang F-J, Roth RH, Wu Y-W, et al. Motor learning selectively strengthens cortical and striatal synapses of motor engram neurons. Neuron. 2022;110(17):2790-2801.e5. doi:10.1016/j.neuron.2022.06.006

8. Sampaio-Baptista C, Scholz J, Jenkinson M, et al. Gray matter volume is associated with rate of subsequent skill learning after a long term training intervention. Neuroimage. 2014;96:158–166. doi:https://doi.org/10.1016/j.neuroimage.2014.03.056

9. Gregory MD, Agam Y, Selvadurai C, et al. Resting state connectivity immediately following learning correlates with subsequent sleep-dependent enhancement of motor task performance. Neuroimage. 2014;102 Pt 2(0 2):666–673. doi:10.1016/j.neuroimage.2014.08.044

10. Sampaio-Baptista C, Filippini N, Stagg CJ, Near J, Scholz J, Johansen-Berg H. Changes in functional connectivity and GABA levels with long-term motor learning. Neuroimage. 2015;106:15–20. doi:10.1016/j.neuroimage.2014.11.032

11. Maruyama S, Fukunaga M, Sugawara SK, Hamano YH, Yamamoto T, Sadato N. Cognitive control affects motor learning through local variations in GABA within the primary motor cortex. Sci Rep. 2021;11(1):18566. doi:10.1038/s41598-021-97974-1

12. Fang Z, Smith DM, Albouy G, et al. Differential Effects of a Nap on Motor Sequence Learning-Related Functional Connectivity Between Young and Older Adults. Front Aging Neurosci. 2021;13. https://www.frontiersin.org/articles/10.3389/fnagi.2021.747358

13. Genzel L, Robertson EM. To Replay, Perchance to Consolidate. PLoS Biol. 2015;13(10):e1002285. doi:10.1371/journal.pbio.1002285

14. Robertson EM, Genzel L. Memories replayed: reactivating past successes and new dilemmas. Philos Trans R Soc B Biol Sci. 2020;375(1799):20190226. doi:10.1098/rstb.2019.0226

15. Ramanathan DS, Gulati T, Ganguly K. Sleep-Dependent Reactivation of Ensembles in Motor Cortex Promotes Skill Consolidation. PLoS Biol. 2015;13(9):e1002263. doi:10.1371/journal.pbio.1002263

16. Tambini A, Davachi L. Awake Reactivation of Prior Experiences Consolidates Memories and Biases Cognition. Trends Cogn Sci. 2019;23(10):876–890. doi:https://doi.org/10.1016/j.tics.2019.07.008

17. Sohal VS, Rubenstein JLR. Excitation-inhibition balance as a framework for investigating mechanisms in neuropsychiatric disorders. Mol Psychiatry. 2019;24(9):1248–1257. doi:10.1038/s41380-019-0426-0

18. Runge K, Cardoso C, de Chevigny A. Dendritic Spine Plasticity: Function and Mechanisms. Front Synaptic Neurosci. 2020;12. https://www.frontiersin.org/articles/10.3389/fnsyn.2020.00036

19. Kolasinski J, Hinson EL, Divanbeighi Zand AP, Rizov A, Emir UE, Stagg CJ. The dynamics of cortical GABA in human motor learning. J Physiol. 2019;597(1):271–282. doi:https://doi.org/10.1113/JP276626

20. Kida H, Mitsushima D. Mechanisms of motor learning mediated by synaptic plasticity in rat primary motor cortex. Neurosci Res. 2018;128:14–18. doi:https://doi.org/10.1016/j.neures.2017.09.008

21. Trepel C, Racine RJ. GABAergic modulation of neocortical long-term potentiation in the freely moving rat. Synapse. 2000;35(2):120–128. doi:10.1002/(SICI)1098-2396(200002)35:2<120::AID-SYN4>3.0.CO;2-6

22. Yasen AL, Smith J, Christie AD. Reliability of glutamate and GABA quantification using proton magnetic resonance spectroscopy. Neurosci Lett. 2017;643:121–124. doi:https://doi.org/10.1016/j.neulet.2017.02.039

23. Stagg CJ, Bachtiar V, Johansen-Berg H. The role of GABA in human motor learning. Curr Biol. 2011;21(6):480–484. doi:10.1016/j.cub.2011.01.069

24. Floyer-Lea A, Wylezinska M, Kincses T, Matthews PM. Rapid modulation of GABA concentration in human sensorimotor cortex during motor learning. J Neurophysiol. 2006;95(3):1639–1644. doi:10.1152/jn.00346.2005

25. Stanley JA, Burgess A, Khatib D, et al. Functional dynamics of hippocampal glutamate during associative learning assessed with in vivo (1)H functional magnetic resonance spectroscopy. Neuroimage. 2017;153:189–197. doi:10.1016/j.neuroimage.2017.03.051

26. Dudai Y, Karni A, Born J. The Consolidation and Transformation of Memory. Neuron. 2015;88(1):20–32. doi:10.1016/j.neuron.2015.09.004

27. Sami S, Robertson EM, Miall RC. The Time Course of Task-Specific Memory Consolidation Effects in Resting State Networks. J Neurosci. 2014;34(11):3982 LP – 3992. doi:10.1523/JNEUROSCI.4341-13.2014

28. Tambini A, Davachi L. Persistence of hippocampal multivoxel patterns into postencoding rest is related to memory. Proc Natl Acad Sci. 2013;110(48):19591 LP –19596. doi:10.1073/pnas.1308499110

29. Doyon J, Benali H. Reorganization and plasticity in the adult brain during learning of motor skills. Curr Opin Neurobiol. 2005;15(2):161–167. doi:10.1016/j.conb.2005.03.004

30. Yousry TA, Schmid UD, Alkadhi H, et al. Localization of the motor hand area to a knob on the precentral gyrus. A new landmark. Brain. 1997;120 (Pt 1:141–157. doi:10.1093/brain/120.1.141

31. Caulo M, Briganti C, Mattei PA, et al. New morphologic variants of the hand motor cortex as seen with MR imaging in a large study population. AJNR Am J Neuroradiol. 2007;28(8):1480–1485. doi:10.3174/ajnr.A0597

32. Kreis R. Issues of spectral quality in clinical 1H-magnetic resonance spectroscopy and a gallery of artifacts. NMR Biomed. 2004;17(6):361–381. doi:10.1002/nbm.891

33. T Vu A, Jamison K, Glasser MF, et al. Tradeoffs in pushing the spatial resolution of fMRI for the 7T Human Connectome Project. Neuroimage. 2017;154:23–32. doi:10.1016/j.neuroimage.2016.11.049

34. Mikkelsen M, Tapper S, Near J, Mostofsky SH, Puts NAJ, Edden RAE. Correcting frequency and phase offsets in MRS data using robust spectral registration. NMR Biomed. 2020;33(10):e4368. doi:https://doi.org/10.1002/nbm.4368

35. Provencher SW. Automatic quantitation of localized in vivo 1H spectra with LCModel. NMR Biomed. 2001;14(4):260–264. doi:https://doi.org/10.1002/nbm.698

36. Near J, Harris AD, Juchem C, et al. Preprocessing, analysis and quantification in single-voxel magnetic resonance spectroscopy: experts’ consensus recommendations. NMR Biomed. 2021;34(5):e4257. doi:10.1002/nbm.4257

37. Gussew A, Erdtel M, Hiepe P, Rzanny R, Reichenbach JR. Absolute quantitation of brain metabolites with respect to heterogeneous tissue compositions in 1H-MR spectroscopic volumes. Magn Reson Mater Physics, Biol Med. 2012;25(5):321–333. doi:10.1007/s10334-012-0305-z

38. Greve DN, Fischl B. Accurate and robust brain image alignment using boundary-based registration. Neuroimage. 2009;48(1):63–72. doi:10.1016/j.neuroimage.2009.06.060

39. Jenkinson M, Smith S. A global optimisation method for robust affine registration of brain images. Med Image Anal. 2001;5(2):143–156. doi:10.1016/s1361-8415(01)00036-6

40. Jenkinson M, Bannister P, Brady M, Smith S. Improved optimization for the robust and accurate linear registration and motion correction of brain images. Neuroimage. 2002;17(2):825–841. doi:10.1016/s1053-8119(02)91132-8

41. Smith SM. Fast robust automated brain extraction. Hum Brain Mapp. 2002;17(3):143–155. doi:10.1002/hbm.10062

42. Dimsdale-Zucker HR, Ranganath C. Chapter 27 - Representational Similarity Analyses: A Practical Guide for Functional MRI Applications. In: Manahan-Vaughan DBT-H of BN, ed. Handbook of Neural Plasticity Techniques. Vol 28. Elsevier; 2018:509–525. doi:https://doi.org/10.1016/B978-0-12-812028-6.00027-6

43. Beckmann CF, Smith SM. Probabilistic independent component analysis for functional magnetic resonance imaging. IEEE Trans Med Imaging. 2004;23(2):137–152. doi:10.1109/TMI.2003.822821

44. Pruim RHR, Mennes M, van Rooij D, Llera A, Buitelaar JK, Beckmann CF. ICA-AROMA: A robust ICA-based strategy for removing motion artifacts from fMRI data. Neuroimage. 2015;112:267–277. doi:10.1016/j.neuroimage.2015.02.064

45. Pruim RHR, Mennes M, Buitelaar JK, Beckmann CF. Evaluation of ICA-AROMA and alternative strategies for motion artifact removal in resting state fMRI. Neuroimage. 2015;112:278–287. doi:10.1016/j.neuroimage.2015.02.063

46. Woolrich MW, Ripley BD, Brady M, Smith SM. Temporal autocorrelation in univariate linear modeling of FMRI data. Neuroimage. 2001;14(6):1370–1386. doi:10.1006/nimg.2001.0931

47. Beckmann CF, Jenkinson M, Smith SM. General multilevel linear modeling for group analysis in FMRI. Neuroimage. 2003;20(2):1052–1063. doi:10.1016/S1053-8119(03)00435-X

48. Woolrich MW, Behrens TEJ, Beckmann CF, Jenkinson M, Smith SM. Multilevel linear modelling for FMRI group analysis using Bayesian inference. Neuroimage. 2004;21(4):1732–1747. doi:10.1016/j.neuroimage.2003.12.023

49. Woolrich M. Robust group analysis using outlier inference. Neuroimage. 2008;41(2):286–301. doi:10.1016/j.neuroimage.2008.02.042

50. Eklund A, Nichols TE, Knutsson H. Cluster failure: Why fMRI inferences for spatial extent have inflated false-positive rates. Proc Natl Acad Sci. 2016;113(28):7900–7905. doi:10.1073/pnas.1602413113

51. Witham CL, Fisher KM, Edgley SA, Baker SN. Corticospinal Inputs to Primate Motoneurons Innervating the Forelimb from Two Divisions of Primary Motor Cortex and Area 3a. J Neurosci. 2016;36(9):2605 LP – 2616. doi:10.1523/JNEUROSCI.4055-15.2016

52. Dubbioso R, Madsen KH, Thielscher A, Siebner HR. The Myelin Content of the Human Precentral Hand Knob Reflects Interindividual Differences in Manual Motor Control at the Physiological and Behavioral Level. J Neurosci. 2021;41(14):3163 LP – 3179. doi:10.1523/JNEUROSCI.0390-20.2021

53. Rathelot J-A, Strick PL. Subdivisions of primary motor cortex based on cortico-motoneuronal cells. Proc Natl Acad Sci. 2009;106(3):918–923. doi:10.1073/pnas.0808362106

54. Josselyn SA, Köhler S, Frankland PW. Finding the engram. Nat Rev Neurosci. 2015;16(9):521–534. doi:10.1038/nrn4000

55. Josselyn SA, Tonegawa S. Memory engrams: Recalling the past and imagining the future. Science. 2020;367(6473). doi:10.1126/science.aaw4325

56. Deshpande G, LaConte S, Peltier S, Hu X. Integrated local correlation: A new measure of local coherence in fMRI data. Hum Brain Mapp. 2009;30(1):13–23. doi:https://doi.org/10.1002/hbm.20482

57. Eisenstein T, Giladi N, Hendler T, Havakuk O, Lerner Y. Physically active lifestyle is associated with attenuation of hippocampal dysfunction in healthy older adults. Front Aging Neurosci. 2021;13(1573):580. doi:10.3389/fnagi.2021.720990

58. Mondragón JD, Marapin R, De Deyn PP, Maurits N. Short- and Long-Term Functional Connectivity Differences Associated with Alzheimer’s Disease Progression. Dement Geriatr Cogn Dis Extra. 2021;11(3):235–249. doi:10.1159/000518233

59. Rajagopalan V, Pioro EP. Disparate voxel based morphometry (VBM) results between SPM and FSL softwares in ALS patients with frontotemporal dementia: which VBM results to consider? BMC Neurol. 2015;15:32. doi:10.1186/s12883-015-0274-8

60. Ashburner J, Friston KJ. Voxel-based morphometry--the methods. Neuroimage. 2000;11(6 Pt 1):805–821. doi:10.1006/nimg.2000.0582

61. Good CD, Johnsrude IS, Ashburner J, Henson RNA, Friston KJ, Frackowiak RSJ. A Voxel-Based Morphometric Study of Ageing in 465 Normal Adult Human Brains. Neuroimage. 2001;14(1):21–36. doi:https://doi.org/10.1006/nimg.2001.0786

62. Yu Z, Guindani M, Grieco SF, Chen L, Holmes TC, Xu X. Beyond t test and ANOVA: applications of mixed-effects models for more rigorous statistical analysis in neuroscience research. Neuron. 2022;110(1):21–35. doi:10.1016/j.neuron.2021.10.030

63. Benjamini Y, Hochberg Y. Controlling the False Discovery Rate: A Practical and Powerful Approach to Multiple Testing. J R Stat Soc Ser B. 1995;57(1):289–300. http://www.jstor.org/stable/2346101

64. Benjamini Y, Yekutieli D. The control of the false discovery rate in multiple testing under dependency. Ann Stat. Published online 2001. doi:10.1214/aos/1013699998

65. Finkelman T, Furman-Haran E, Paz R, Tal A. Quantifying the excitatory-inhibitory balance: A comparison of SemiLASER and MEGA-SemiLASER for simultaneously measuring GABA and glutamate at 7T. Neuroimage. 2022;247:118810. doi:https://doi.org/10.1016/j.neuroimage.2021.118810

66. Press DZ, Casement MD, Pascual-Leone A, Robertson EM. The time course of off-line motor sequence learning. Cogn Brain Res. 2005;25(1):375–378. doi:https://doi.org/10.1016/j.cogbrainres.2005.05.010

67. Rideaux R. No balance between glutamate+glutamine and GABA+ in visual or motor cortices of the human brain: A magnetic resonance spectroscopy study. Neuroimage. 2021;237:118191. doi:https://doi.org/10.1016/j.neuroimage.2021.118191

68. Rubin R, Abbott LF, Sompolinsky H. Balanced excitation and inhibition are required for high-capacity, noise-robust neuronal selectivity. Proc Natl Acad Sci. 2017;114(44):E9366–E9375. doi:10.1073/pnas.1705841114

69. Zhou S, Yu Y. Synaptic E-I Balance Underlies Efficient Neural Coding. Front Neurosci. 2018;12. https://www.frontiersin.org/articles/10.3389/fnins.2018.00046

70. Lim S, Goldman MS. Balanced cortical microcircuitry for maintaining information in working memory. Nat Neurosci. 2013;16(9):1306–1314. doi:10.1038/nn.3492

71. Feldman DE. Synaptic mechanisms for plasticity in neocortex. Annu Rev Neurosci. 2009;32:33–55. doi:10.1146/annurev.neuro.051508.135516

72. Rioult-Pedotti MS, Friedman D, Donoghue JP. Learning-induced LTP in neocortex. Science. 2000;290(5491):533–536. doi:10.1126/science.290.5491.533

73. Rioult-Pedotti MS, Friedman D, Hess G, Donoghue JP. Strengthening of horizontal cortical connections following skill learning. Nat Neurosci. 1998;1(3):230–234. doi:10.1038/678

74. Ziemann U, Ilić T V, Pauli C, Meintzschel F, Ruge D. Learning modifies subsequent induction of long-term potentiation-like and long-term depression-like plasticity in human motor cortex. J Neurosci Off J Soc Neurosci. 2004;24(7):1666–1672. doi:10.1523/JNEUROSCI.5016-03.2004

75. Markram H, Tsodyks M. Redistribution of synaptic efficacy between neocortical pyramidal neurons. Nature. 1996;382(6594):807–810. doi:10.1038/382807a0

76. Eder M, Zieglgänsberger W, Dodt H-U. Neocortical long-term potentiation and long-term depression: site of expression investigated by infrared-guided laser stimulation. J Neurosci Off J Soc Neurosci. 2002;22(17):7558–7568. doi:10.1523/JNEUROSCI.22-17-07558.2002

77. Zakharenko SS, Zablow L, Siegelbaum SA. Visualization of changes in presynaptic function during long-term synaptic plasticity. Nat Neurosci. 2001;4(7):711–717. doi:10.1038/89498

78. Lea-Carnall CA, El-Deredy W, Williams SR, Stagg CJ, Trujillo-Barreto NJ. From synaptic activity to human in vivo quantification of neurotransmitter dynamics: a neural modelling approach. bioRxiv. Published online January 1, 2021:2021.03.11.434540. doi:10.1101/2021.03.11.434540

79. Doyon J, Gabitov E, Vahdat S, Lungu O, Boutin A. Current issues related to motor sequence learning in humans. Curr Opin Behav Sci. 2018;20:89–97. doi:https://doi.org/10.1016/j.cobeha.2017.11.012

80. Carr MF, Jadhav SP, Frank LM. Hippocampal replay in the awake state: a potential substrate for memory consolidation and retrieval. Nat Neurosci. 2011;14(2):147–153. doi:10.1038/nn.2732

81. Castro-Alamancos MA, Donoghue JP, Connors BW. Different forms of synaptic plasticity in somatosensory and motor areas of the neocortex. J Neurosci. 1995;15(7):5324 LP – 5333. doi:10.1523/JNEUROSCI.15-07-05324.1995

82. Chen SX, Kim AN, Peters AJ, Komiyama T. Subtype-specific plasticity of inhibitory circuits in motor cortex during motor learning. Nat Neurosci. 2015;18(8):1109–1115. doi:10.1038/nn.4049

83. Stagg CJ, Bachtiar V, Amadi U, et al. Local GABA concentration is related to network-level resting functional connectivity. Culham JC, ed. Elife. 2014;3:e01465. doi:10.7554/eLife.01465

84. Noguchi J, Nagaoka A, Hayama T, et al. Bidirectional in vivo structural dendritic spine plasticity revealed by two-photon glutamate uncaging in the mouse neocortex. Sci Rep. 2019;9(1):13922. doi:10.1038/s41598-019-50445-0

85. Draganski B, Gaser C, Busch V, Schuierer G, Bogdahn U, May A. Changes in grey matter induced by training. Nature. 2004;427(6972):311–312. doi:10.1038/427311a

86. Taubert M, Draganski B, Anwander A, et al. Dynamic properties of human brain structure: learning-related changes in cortical areas and associated fiber connections. J Neurosci Off J Soc Neurosci. 2010;30(35):11670–11677. doi:10.1523/JNEUROSCI.2567-10.2010

87. Markram H, Toledo-Rodriguez M, Wang Y, Gupta A, Silberberg G, Wu C. Interneurons of the neocortical inhibitory system. Nat Rev Neurosci. 2004;5(10):793–807. doi:10.1038/nrn1519

88. Stagg CJ. Magnetic Resonance Spectroscopy as a tool to study the role of GABA in motor-cortical plasticity. Neuroimage. 2014;86:19–27. doi:10.1016/j.neuroimage.2013.01.009

89. Choi C-H, Iordanishvili E, Shah NJ, Binkofski F. Magnetic resonance spectroscopy with transcranial direct current stimulation to explore the underlying biochemical and physiological mechanism of the human brain: A systematic review. Hum Brain Mapp. 2021;42(8):2642–2671. doi:10.1002/hbm.25388

