## Supplementary Materials for "Early excitatory and inhibitory modifications in the motor cortex following skill learning support motor memory consolidation and cortical plasticity overnight"

### Supplementary material

#### MRS quality metrics

The MRS quality metrics are presented in **Table S1**. Out of 216 MRS measurements across all participants (36 participants x 6 MRS runs), 5 measurements (across 3 participants) were not included in the final data analyses due to low SNR (i.e., <30) or contaminated spectrum, and 2 other measurements from an additional participant were not acquired due to time limitation in the scanner. Repeated measures mixed model analysis confirmed no significant differences in the voxels GM fraction ($p$=.601), WM fraction ($p$=.507), or CSF fraction ($p$=.560) between the two sessions. Furthermore, water linewidth was not statistically different between each pair of MRS runs over the two sessions ($p$>.05, FDR-corrected). The SNR on the second day MRS run was significantly higher compared to each post-learning run on day 1 ($p$<.05, FDR-corrected). No differences were found between each two runs within day 1 ($p$>.19 for all pairwise comparisons, FDR-corrected). **Figure S1** demonstrates a representative spectrum acquired from one participant.

**Table S1.** MRS quality metrics (Mean±SD)

| MRS run | Water FWHM | SNR | CRLB (GABA) | Voxel GM fraction | Voxel WM fraction | Voxel CSF fraction |
|---|---|---|---|---|---|---|
| Pre-learning | 10.67±1.18 | 59.06±5.67 | 29.33±8.3 | 0.373±0.03 | 0.575±0.05 | 0.052±0.03 |
| Post 5 min | 10.79±1.08 | 58.83±5.68 | 33.54±15.3 | 0.373±0.03 | 0.575±0.05 | 0.052±0.03 |
| Post 20 min | 10.67±1.09 | 58.49±5.47 | 28.51±7.2 | 0.373±0.03 | 0.575±0.05 | 0.052±0.03 |
| Post 25 min | 10.81±1.22 | 57.28±6.14 | 32.90±12.6 | 0.373±0.03 | 0.575±0.05 | 0.052±0.03 |
| Post 30 min | 10.94±1.18 | 58.1±5.11 | 29.06±5.9 | 0.373±0.03 | 0.575±0.05 | 0.052±0.03 |
| Overnight | 10.58±1.09 | 60.83±5.78 | 29.86±9.7 | 0.370±0.03 | 0.579±0.05 | 0.050±0.03 |

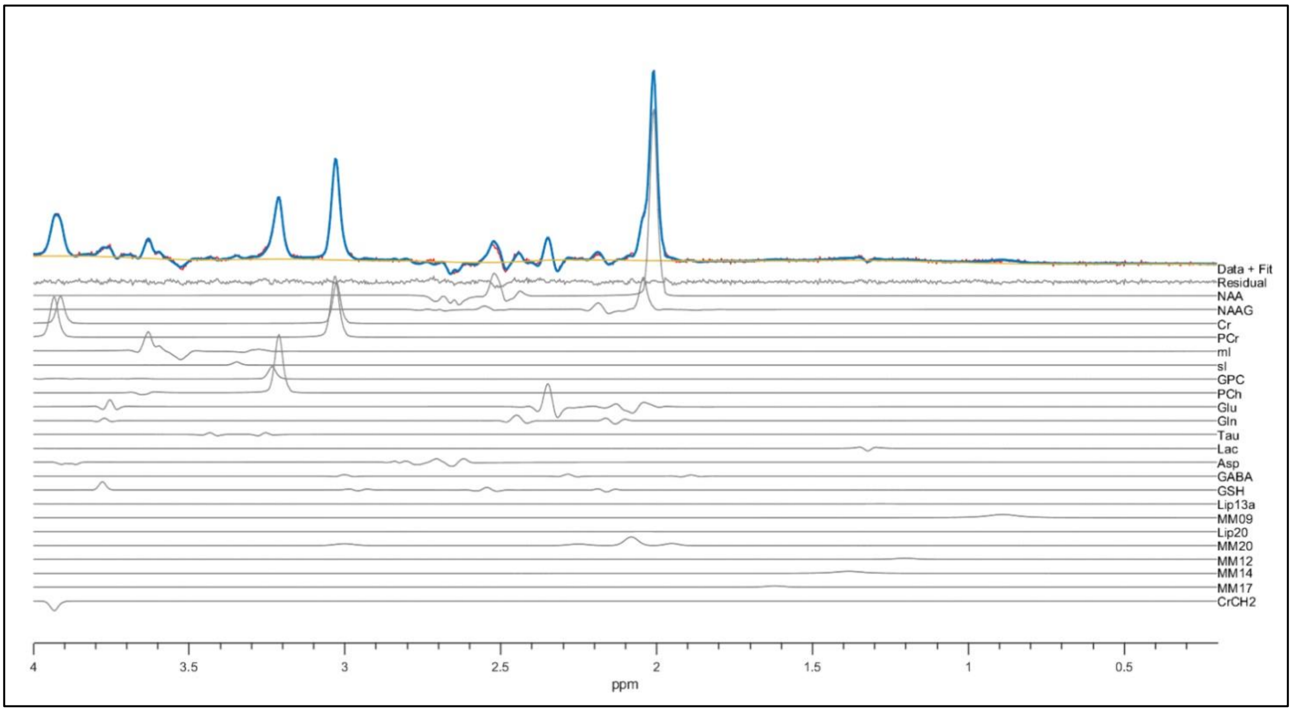


**Figure 1.** A representative spectrum acquisition from one participant including model fit.
